# Confocal and Transmission electron microscopy imaging of *Orientia tsutsugamushi*

**DOI:** 10.64898/2026.08.24.746599

**Authors:** Meenakshi Rana, Suvradeep Mitra, Mohan Kumar Hanumanthappa, Navneet Sharma, Manisha Biswal

## Abstract

Scrub typhus, caused by *Orientia tsutsugamushi*, is an obligate intracellular gram-negative pathogen that remains a cause of acute febrile illness in India. Culture isolation of *Orientia tsutsugamushi* clinical isolates is infrequent because it is technically more challenging than PCR-based molecular identification.

In this report, we describe the culture isolation of *Orientia* from the whole blood of a 64-year-old farmer with acute febrile illness. Whole blood was inoculated onto an 80% confluent L929 cell line. Real-time PCR targeting the 47-kDa and 56-kDa genes, combined with Sanger sequencing, confirmed the isolate. Transmission electron microscopy of infected L929 cells revealed multiple oval-shaped bacteria within the host cytoplasm. Confocal microscopy demonstrated progressive accumulation of CFSE-labelled bacteria within infected cells over time. These findings support the successful isolation and visualization of a blood-derived *O. tsutsugamushi* isolate and provide a platform for downstream assays of host-pathogen interactions, antimicrobial susceptibility testing, and vaccine development.

## Introduction

Scrub typhus, a major cause of acute febrile illness in India, is caused by *Orientia tsutsugamushi*, which is a gram-negative obligate intracellular bacterium ^[1]^. The bacterium can only replicate within living host cells; its isolation requires specialized cell culture techniques making the isolation challenging. The diagnosis and molecular epidemiology studies rely entirely on PCR-based assays. Successful culture of *Orientia* is important for studying bacterial biology and antigenic variation which are important for designing appropriate diagnostics and vaccines. Growth of *Orientia tsutsugamushi* from patient blood samples has been attempted in Vero cell lines in South India ^[2]^. Isolates from different geographical locations of a vast country like India is required for country-wide representative data. Here, we report the isolation of *Orientia tsutsugamushi* from infected human blood collected in north India and present microscopic visualization of the isolate using confocal microscopy and transmission electron microscopy (TEM).

### Case report

A 64-year-old farmer from Bilaspur, Himachal Pradesh, a state in north India, was admitted to the emergency department in the first week of October,2024 with four days of fever, thrombocytopenia and shock. The patient had received one dose of doxycycline at the referring hospital. History revealed exposure to vegetation while collecting fodder for cattle.

On examination, the patient had no eschar. Investigations showed a platelet count of 12 × 10^9^/L (normal range:150× 10^9^/L – 450× 10^9^/L) and a hemoglobin of 12.3 g/dL (normal range: 13.5g/dL – 17.5g/dL). The rapid IgM serological test against *Orientia tsutsugamushi* was positive at bedside. Samples were collected for dengue serology. Blood was collected prior to any antibiotic treatment for PCR and culture of scrub typhus. The patient was started on doxycycline and resuscitated with fluids. He improved over the next seven days and was discharged.

### Investigations for scrub typhus

DNA was isolated from blood using the Qiagen DNA extraction kit and was subjected to 47-kDa real-time PCR, which was positive (Ct value 18)^[3]^. The genotype of the strain was confirmed to be Karp-like (GenBank accession number PZ023951) using 56-kDa nested PCR and subsequent Sanger sequencing ^4]^.

Within 1 hour of sample collection, whole blood was diluted with an equal volume of 1X PBS and inoculated onto 80% confluent L929 cells in a 6-well plate in duplicate. The plate was incubated for 30 minutes at 37°Celsius in a 5% CO_2_ incubator with gentle agitation at regular intervals. The diluted blood was removed from the wells and washed two times with 1X PBS to remove any residual blood. The wells were then filled with DMEM containing 10% FBS. Media were changed every two days. Cells were cultured for seven days. After 7 days, the cells were passaged and checked for signs of contamination and morphological changes until the next passage, with media changes every 2 days. After 14 days, the cells were subjected to real-time PCR targeting the 47-kDa gene, which was positive (Ct value 28). The positive culture was maintained in T25 flasks until the cell load was sufficient (Ct value 12) for TEM and confocal imaging.

### Transmission electron microscopy (TEM) imaging

The cell scrapings from positive culture flask were subjected to centrifugation at 1000 rpm for 15 minutes and were stored in 0.2% glutaraldehyde at 4°C for pre-fixing. The cells were again pelleted down at 3000 rpm for five minutes. The pellet was then washed with two changes of Sorenson’s buffer with sucrose, post-fixed with 1% osmic acid in Millonig’s buffer and incubated at 4°C for two hours. After two hours the cells were washed with two changes of Millonig’s buffer. Next, the cells were dehydrated using a series of ascending alcohol concentrations (70%-100%) followed by two changes of propylene oxide treatment to replace the alcohol in cells and help the penetration of epoxy mix into the cells. Next, the cells were embedded with epoxy resin on rubber molds with dimethyaminomethyl phenol (DMP-30) as accelerator and kept for polymerization at 60°C overnight. One-micron (1µ) thick sections were cut to select the area for ultrathin sections, which were cut at a setting of 70nm, using ultracut and mounted on copper 200 mesh grids. The sections were stained with uranyl acetate and lead citrate by standard technique. Grids were viewed using JEOL JEM 1400 transmission electron microscope. Photographs were taken under fixed magnifications from appropriate areas.

### Confocal Imaging

The 80% confluent L929 cell lines were infected with the isolated *Orientia* and incubated for seven days. The bacteria were purified from L929 cell lines using the protocol published by Giengkam et al ^[5]^. In brief, the cells were scraped from the flasks and centrifuged at 4000 rpm for 10 minutes. The host cells were lysed using 0.2-mm glass beads and pulse vortexed.

The lysate was centrifuged at 4000 rpm for 5 minutes to pellet lysed host cells and their debris. The supernatant was collected and ultracentrifuged for 5 minutes to pellet the bacteria. The pelleted bacteria were then used for further processing.

The pelleted bacteria were then suspended in 5 μM of CFSE dissolved in DMSO and incubated for 15 minutes at 37°C in the dark. The bacteria were pelleted again, suspended in DMEM with 5% FBS, and incubated at room temperature for 30 minutes to remove excess dye. Bacteria were pelleted again and suspended in fresh DMEM containing 5% FBS. The bacteria were then added to 80% confluent L929 cell lines in 6-well plates ^[6]^. The cells were then fixed with 4% paraformaldehyde at 8, 16, and 24 hours. The fixed cells were then stained with 1µg/mL DAPI and visualised under a confocal microscope.

## Results

### Transmission electron microscopy reveals abundant intracellular *O. tsutsugamushi* in infected L929 monolayer

Transmission electron microscopy of infected L929 cells revealed multiple oval-shaped bacteria consistent with *Orientia tsutsugamushi* (marked with red stars). These round-shaped bacteria (856-nm in size; Figure 1(A)) were observed within the cytoplasm of a lysed host cell. The other large organelles, such as mitochondria, were identified by the presence of internal cristae (marked with blue stars) (Figure 1).

**Figure 1.**
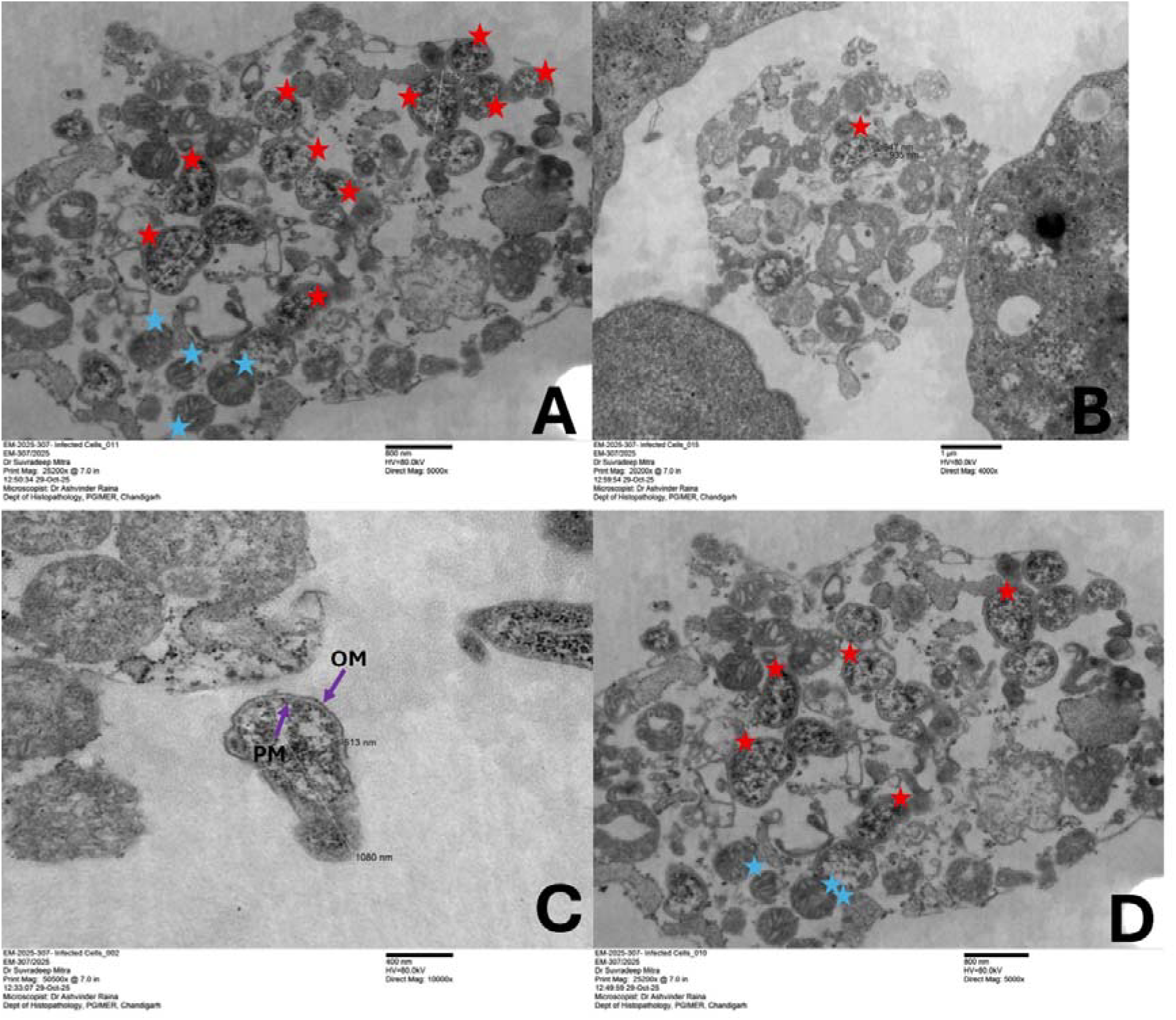
Transmission electron microscopy revealing intracellular *Orientia tsutsugamushi* in infected L929 cell lines, with representative images and size measurements. Low-magnification images (panels A and B) display multiple bacteria consistent with *Orientia* (red stars) within host cytoplasm; host mitochondria with characteristic cristae (blue stars); host nucleus (yellow lines). A high-magnification image (10,000×) shows the measured bacterial dimensions of 513 nm × 1080 nm. Scale bars are provided in each panel.

### Time-dependent intracellular accumulation of CFSE-labelled *O. tsutsugamushi* in L929 cell culture

Intracellular localisation of CFSE-labelled *Orientia tsutsugamushi* within L929 cell lines was also demonstrated using confocal imaging. Control L929 cell lines displayed minimal background fluorescence in the green channel. In infected cells, CFSE-positive fluorescence was observed in the cytoplasm of infected L929 cells as early as 8 hours post-infection, consistent with bacterial uptake or entry into the fibroblast cell lines. By 16 hours, the green fluorescence signal was frequently detected throughout the cells. At 24 hours, there was an increased CFSE-labelled fluorescence throughout the infected monolayer consistent with bacterial accumulation over time. A few extracellular signals were also observed after 16 hours, reflecting progressive intracellular bacterial accumulation, followed by host-cell membrane damage, partial lysis, and the release of labelled bacteria into the extracellular milieu (Figure 2).

**Figure 2.**
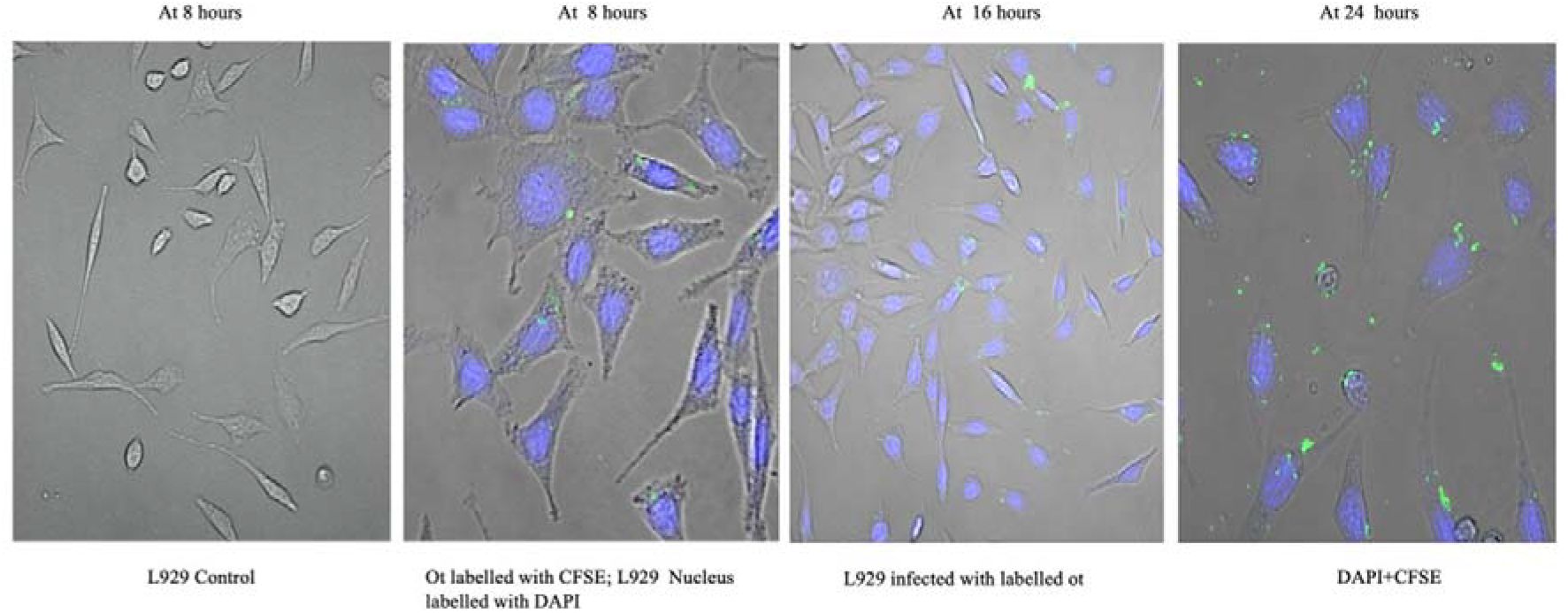
Confocal microscopy demonstrating intracellular accumulation of Orientia tsutsugamushi in L929 cell lines. Panel 1 (left): Uninfected L929 control cells showing minimal background green signal. Panels 2-4: L929 cells infected with CFSE-labelled O. tsutsugamushi at 8 hours, 16 hours, and 24 hours post-infection, showing progressive green fluorescence (CFSE signal) within the cytoplasm. Host cell nuclei are counterstained with DAPI (blue). Progressive intensification of the green signal demonstrates time-dependent bacterial accumulation and multiplication within infected cells.

## Discussion

In the present study, *Orientia* was isolated from the blood of a patient with acute febrile illness. *Orientia* isolation was confirmed from a PCR-positive blood sample in an in vitro cell culture system. This is the first report of isolation of *O. tsutsugamushi* from any clinical specimen from Northern India.

Isolation of this bacterium and its successful maintenance in cell lines offer many avenues of research on this neglected intracellular pathogen. Blood-derived clinical isolates may exhibit phenotypic differences compared with lab-adapted strains, including ATCC reference strains. Continuous passaging and long-term storage select for variants better suited to cell culture, which may not accurately reflect in vivo infection dynamics. Scrub typhus is a huge public health problem in India^[7]^. Multiple circulating strains of *Orientia* exist across endemic regions, showing geographical variation that may reflect strain-specific differences. However, most scrub typhus research has utilised reference strains (Karp, Gilliam, Kato, Boryong, UT76, UT176) isolated decades ago ^[8]^These strains remain important reference models but may not represent the biology of currently circulating clinical isolates. India-specific strains, freshly isolated from patients, would be invaluable for conducting studies on diagnostics, pathogenesis, and vaccine development in the Indian context. Recent reports from Southern India describe isolation of clinical isolates from human peripheral blood mononuclear cells (PBMCs), but limited data are available from North India. The current North India isolate may provide valuable comparative data on regional strain variations.

TEM imaging revealed the ultrastructural features consistent with *O. tsutsugamushi*, small oval-shaped bacteria within host-cell cytoplasm^[9]^.The presence of multiple organisms across multiple fields strengthens the successful isolation of *Orientia*. These findings were consistent with the positive real-time PCR results in infected L929 cells.

Confocal microscopy was successfully standardised in this study. The bacteria showed green fluorescence and multiplication over time, while the control showed negligible green signal, demonstrating the specificity of the signal in the CFSE channel. Intracellular fluorescence was detectable as early as 8 hours, with progressive intensification by 24 hours. The live cell imaging of CFSE-labelled bacteria inside the host cell cytoplasm enables further studies on the pathogenesis of this infection

## Conclusion

We successfully isolated *Orientia tsutsugamushi* from the PCR-confirmed patient’s whole blood using L929 mouse fibroblast cell lines. The isolation was confirmed by imaging techniques. Confocal imaging revealed progressive accumulation of CFSE-labelled fluorescence within the host cell cytoplasm over time. TEM revealed ultrastructural features consistent with *O. tsutsugamushi*. This isolate provides a platform for quantitative antimicrobial susceptibility assays, host-pathogen interaction studies, and vaccine development, addressing critical research gaps in scrub typhus.

## Supporting information

Phylogenetic Tree

## Data availability

The 56-kDa sequences are deposited at NCBI with GenBank accession number PZ023951.

## Ethics Statement

This work is a part of a PhD thesis approved by the Institutional Ethics Committee. The institutional ethics approval number is IEC-INT/2024/PhD-1956.

## Supplementary information

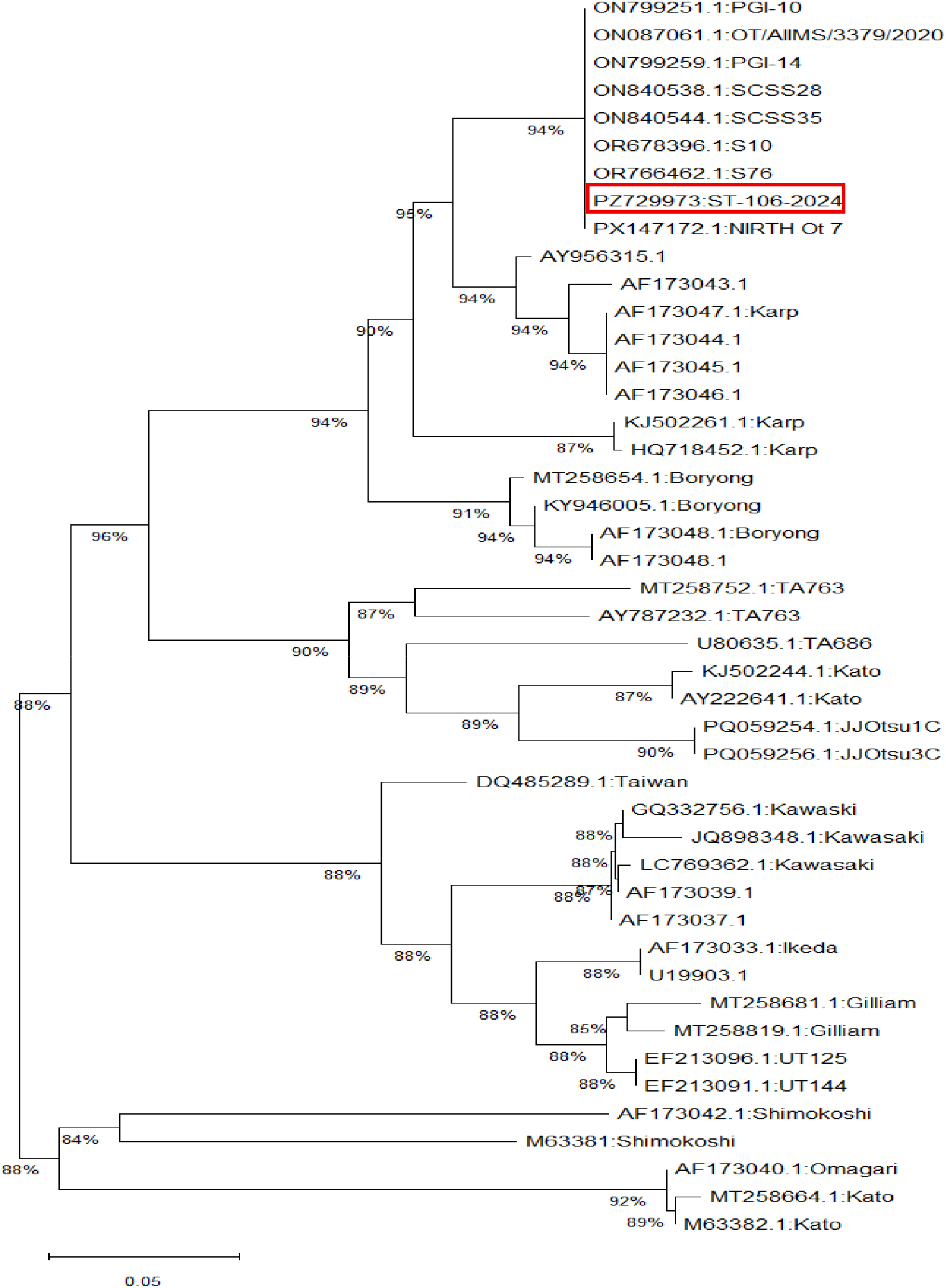
Phylogenetic analysis of *Orientia tsutsugamushi* based on partial 56-kDa type-specific antigen gene sequences. The phylogenetic tree was constructed using the maximum-likelihood method in MEGA 12. Branch support was evaluated using 1,000 bootstrap replicates, and bootstrap values are shown at the nodes. The sequence generated in the present study is indicated by red block. Reference sequences were retrieved from GenBank, and accession numbers are provided alongside the sequence names.

