## Supplementary material for "Confocal and Transmission electron microscopy imaging of *Orientia tsutsugamushi*": Phylogenetic Tree

**Supplementary information:**

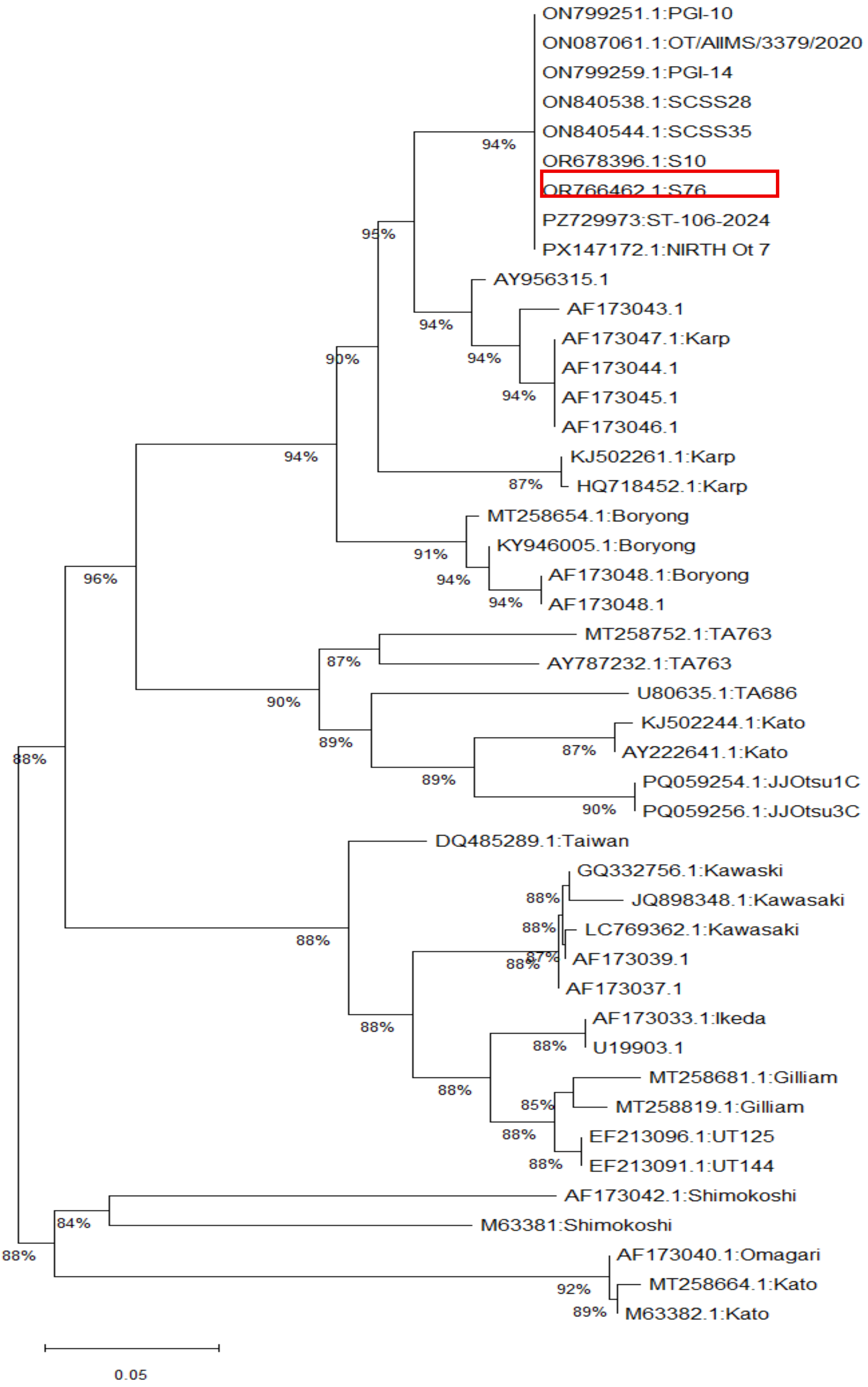

**Phylogenetic analysis of *Orientia tsutsugamushi* based on partial 56-kDa type-specific antigen gene sequences.** The phylogenetic tree was constructed using the maximum-likelihood method in MEGA 12. Branch support was evaluated using 1,000 bootstrap replicates, and bootstrap values are shown at the nodes. The sequence generated in the present study is indicated by red block. Reference sequences were retrieved from GenBank, and accession numbers are provided alongside the sequence names.
